# Energy and time trade-offs explain everyday human reaching movements

**DOI:** 10.1101/2025.05.16.654498

**Authors:** Jeremy D. Wong, William J. Herspiegel, Arthur D. Kuo

**Affiliations:** Dept. of Biomedical Engineering, University of Calgary, Alberta, CANADA; Faculty of Kinesiology, University of Calgary, Alberta, CANADA

**Keywords:** metabolic energetic cost, temporal discounting, upper extremity, smoothness, minimum jerk

## Abstract

Humans perform reaching movements with stereotypically smooth trajectories, attributed to neural optimality principles. Previous models focus on maximizing objectives like accuracy to explain smooth reaching under the speed-accuracy tradeoff, but none explain individuality. In everyday tasks such as reaching for a cup or pen, individuals self-select their own relaxed and idiosyncratic speeds that prioritize neither speed nor accuracy and appear to defy purely objective optimality. Here we propose an Energy-Time trade-off that better predicts the smoothness, speed, and individuality of everyday reaching. Energy refers to metabolic energy expenditure, and Time represents a weighted cost adjustable for individual and contextual factors. The balance between the two predicts human speed trajectories, durations, and peak speeds more accurately than prior models. Individuals move differently because they value time individualistically yet consistently. For example, one’s time valuation may be inferred from a single reach, and then applied to predict reaches of any distance by the same individual. Instructions to reach “faster” or “slower” also yield individualistic yet identifiable time valuations sufficient to predict other movements within each context. In fact, all movements across individuals and contexts align with a universal set of Energy-Time predictions. Energy economy objectively favors smoothness and slowness, whereas the time cost captures one’s individualistic preference to spend energy to save time.

## Introduction

Humans reach for targets with stereotypical motions characterized by smoothness and bell-shaped velocity profiles (Fig. 1A). Point-to-point reaches are typically ballistic and pre-programmed (1), thought to reflect neural optimality principles that resolve motor redundancy. Proposed objectives include maximizing smoothness (2, 3) or, via the speed-accuracy tradeoff, accuracy (4, 5). Movement durations and peak speeds also scale systematically with distance (6–8). However, criteria proposed to date fail to explain everyday reaching tasks—such as grabbing a cup—that demand little speed or accuracy yet remain smooth. Humans can also modulate movement speed based on subjective context (e.g., hurrying), which previous theories do not address. A broader framework is needed to encompass everyday reaching movements.

**Fig. 1.**
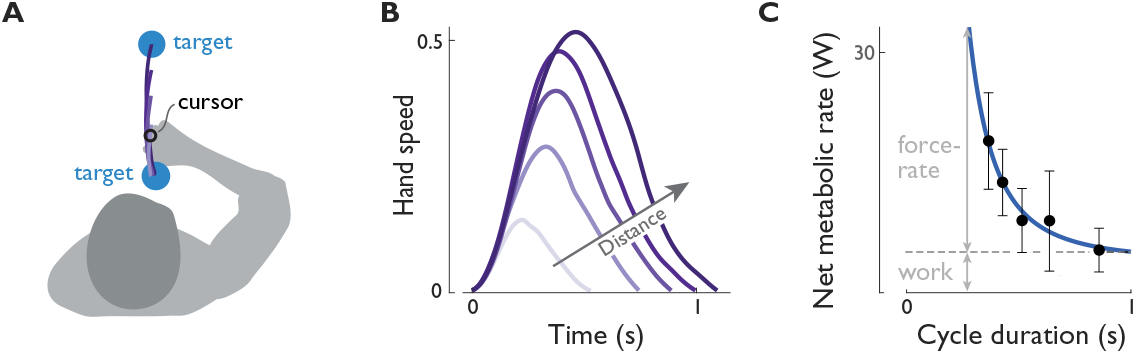
Humans perform (**A**) point-to-point reaching movements with (**B**) smooth, bell-shaped speed trajectories^1^, where preferred speed and duration increasing with distance^2,3^. The speed-accuracy trade-off explains smooth trajectories for fast or accurate movements^4^ but does not account for slower, “everyday” movements of low accuracy or their durations. (**C**) Reaching movements incur increasing metabolic energy costs for shorter durations^5^, as illustrated for cyclic movements of varying duration but fixed work per cycle^6^. Faster movements are energetically costly due to a term called force-rate, which is proportional to the peak rate of force development during a movement. Arm movements are between visually displayed targets and physically supported by a planar two-link manipulandum.

Several objectives have been proposed to explain smooth speed trajectories, starting with minimizing kinematic jerk (third derivative of position 9, 2). Later work incorporated muscle forces as a torque-change objective (3). A separate hypothesis is that larger motor commands are also noisier, which could explain the speed-accuracy tradeoff by linking accuracy to smoothness (4). These objectives are appealing because they can explain various conditions, such as moving against a spring-like load (3) or in figure-8 paths (4). However, most cannot generally explain movement durations. Most are also abstract and rely on hypothesized signals not directly observable in experiments. Their connection to physiology therefore remains elusive.

A similar challenge applies to movement durations and speeds. Reaching movements, like saccadic eye movements (10), exhibit a one-dimensional relationship between movement distance, duration, and peak speed (6). These relationships have been attributed to criteria such as motor noise with a signal-independent component (11), temporally discounted energy or effort (12, 13), or various functions of time (7, 14, 15, 8). Metabolic energy is particularly appealing due to its biological importance for survival (16) and its role in neuronal processes (17). For example, energy expenditure during adaptation to novel reaching dynamics (18) and increases with faster movements (Figure 1B 12, 14). Time is valuable because that spent moving could be spent on other tasks or competing goals. A combination of energy- and time-related objectives could potentially explain movement trajectories and durations using biologically identifiable signals (8, 14, 19).

Most daily movements are undemanding of speed or accuracy. For example, one might unhurriedly reach for a pen or cup based on memory of its location or approximate peripheral vision. Objects that are frequently used tend to be easy to reach. Such reaches occur at a self-paced speeds that are not universal but appear idiosyncratic to the individual (19). Humans also adjust their speed depending on subjective factors like urgency. This modulation could reflect a subjective valuation for time; however, such free parameters often require fitting to data making a given theory difficult to falsify. A viable explanation must account for subjectivity and contextual dependence while also making testable predictions.

We propose an objective for undemanding “Everyday” point-to-point reaching that combines minimizing energy expenditure and time (Energy-Time, Fig. 2). We used a previously developed representation of energy and time costs (14), where the time cost is linearly proportional to a a subjective coefficient, *c*_*T*_. Here we tested whether these terms (Fig. 2A) together can predict movement durations *T*, peak speeds *V*, and hand speed trajectories *ν*(*t)* for any movement distance *D* (Fig. 2B). We also tested whether the subjective time cost could accommodate different individuals and task contexts. This required identifying the time valuation *c*_*T*_, which we used to independently predict other movements within the same context.

**Fig. 2.**
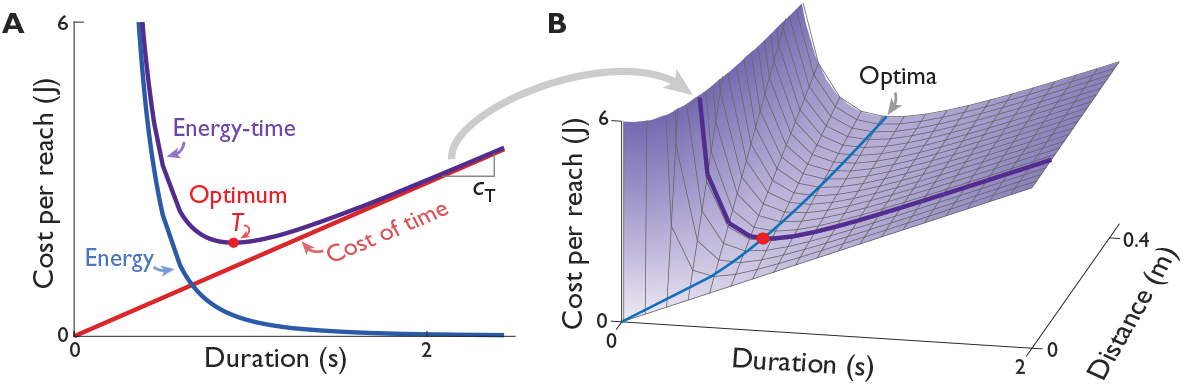
The Energy-Time hypothesis for reaching. (**A**) The objective function consists of two terms: an energy cost (Eq. 1, consistent with data in Fig. 1**C**) and a proportional cost for time. Their sum predicts an optimal movement duration, adjustable by an individual-specific time valuation *c*_*T*_, a slope with units of energy per time. (**B**) The proposed cost objective also depends on movement distance, forming a surface when plotted against duration and distance. The model predicts an optimal duration (labeled Optima) increasing with distance (Eq. 2) for low-accuracy, “Everyday” tasks.

If the tradeoff between energy and time passes these tests, it may provide a more comprehensive explanation for everyday reaching than previous theories.

## Results

### Model predictions

Analysis of the Energy-Time model yields simple predictive power laws. While optimal control is required to predict full movement trajectories, the key predictions for movement duration and peak speed can be summarized by focusing only on the leading terms of the cost function. The approximate objective for moving a mass *M* by distance *D* in duration *T* consists of two main terms:

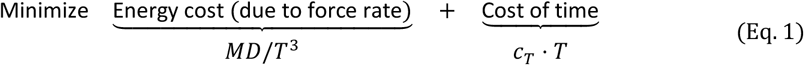

Metabolic energy cost also depends on muscle work but is dominated by force-rate (Fig. 1C), which refers to the peak rate of force development (14). The movement duration *T* is valued proportionally with a coefficient *c*_*T*_ with units of energy per time. Optimizing these two terms yields the following predictions:

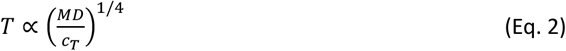

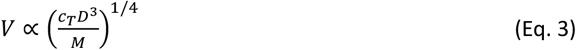

Thus, durations are expected to increase with the one-fourth power of distance, and peak speeds with the three-fourths power. Although hand speed trajectories require full optimal control calculations, these power laws suffice to predict duration (Fig. 2B) and peak speeds using only a single empirical parameter: the time valuation *c*_*T*_.

### Experimental results

#### Everyday reaching predicted from a single data point

The Energy-Time model predicts reaching motions from a single data point (Fig. 3). An individual’s self-selected duration for an Everyday reach was used to estimate their time valuation *c*_*T*_ (Fig. 3**A**–**B**), representing the marginal energy they are willing to expend to save a unit of time. This single parameter then predicted durations *T*, peak speeds *V* (Fig. 3**B**–**C**), and trajectories (Fig. 3**D**) across distances *D* not yet encountered. This approach contrasts with prior models that require multiple parameters or extensive data fitting, highlighting the simplicity and predictive power of the Energy-Time model.

**Fig 3.**
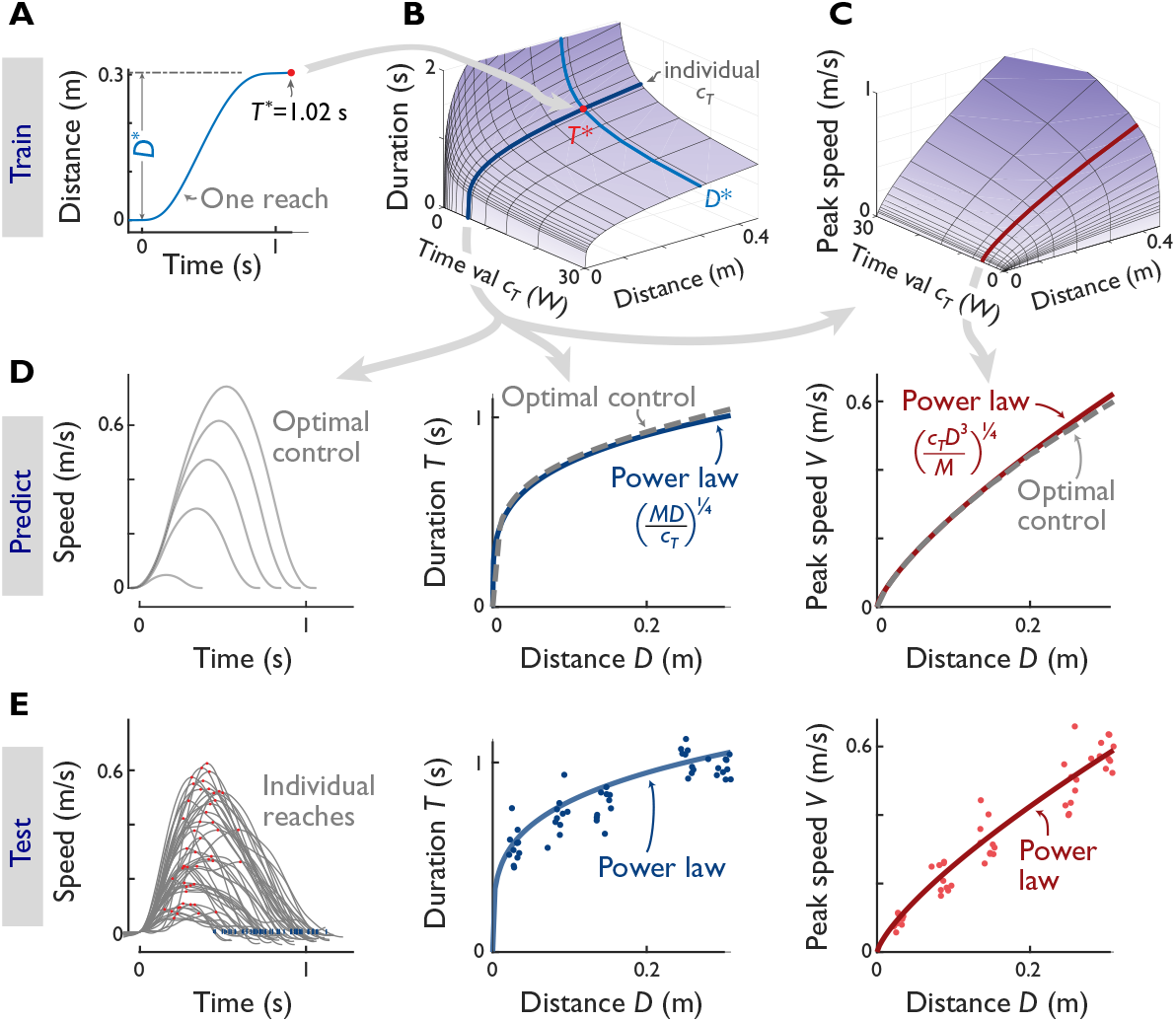
The Energy-Time model uses a single observation to predict reaching behavior for an individual. (**A**) The observation is the individual’s time duration *T*^∗^ (red dot) to reach a fixed distance *D*^∗^ (e.g., 0.29 m in 0.99 s), which determines their time valuation *c*_*T*_. (**B**) The model predicts the relationship between duration, distance, and *c*_*T*_: measuring *T*^∗^ (red dot) for given *D*^∗^ (light blue line) determines individual’s *c*_*T*_ (e.g., 4.0 W), which in turn predicts durations *T* and (**C**) peak speeds *V* for any distance *D* (solid lines for constant *c*_*T*_). (**D**) Optimal control model then predicts how trajectories, durations, and peak speeds vary with distance, for the individual’s specific time valuation. Power law approximations summarize these predictions (*R*^2^ = 0.98 for durations, 1.00 for peak speeds), yielding scalable functions for duration (Eq. 2) and peak speed (Eq. 3). (**E**) Experimental data from the same individual reaching five different targets agree with model predictions (*R*^2^= 0.77 for trajectories, 12.6% RMSE for durations and 18.8% RMSE for peak speeds, root-mean-square error). Results from single observation procedure for all individuals (*N* = 9) are shown in Table 1.

Experiments agreed with these single-observation predictions. For illustration, one individual performed a self-paced Everyday reach of 0.29 m (to a 2.5 cm diameter target) in 0.99 s (Fig. 3A), yielding an estimated *c*_*T*_ of 4.0 W. Predictions using this value were tested with five additional distances, showing strong agreement with experimental data (*R*^2^= 0.77 for trajectories, RMSE = 12.6% for durations and 18.8% for peak speeds, root-mean-square error; Fig. 3**E**). Across all participants (*N* = 9), single-observation estimates yielded similar results (*c*_*T*_ = 3.8 W, *R*^2^ = 0.72; Table 1 Everyday single observation). This single observation per individual therefore predicted bell-shaped speed trajectories, durations, and peak speeds across a range of distances. Moreover, power law predictions (Eqs. 2–3) aligned closely with optimal control results (*R*^2^ = 0.98 for durations and 1.00 for peak speeds; Fig. 3**D**). These power laws are therefore treated as model predictions from this point forward.

**Table 1.**
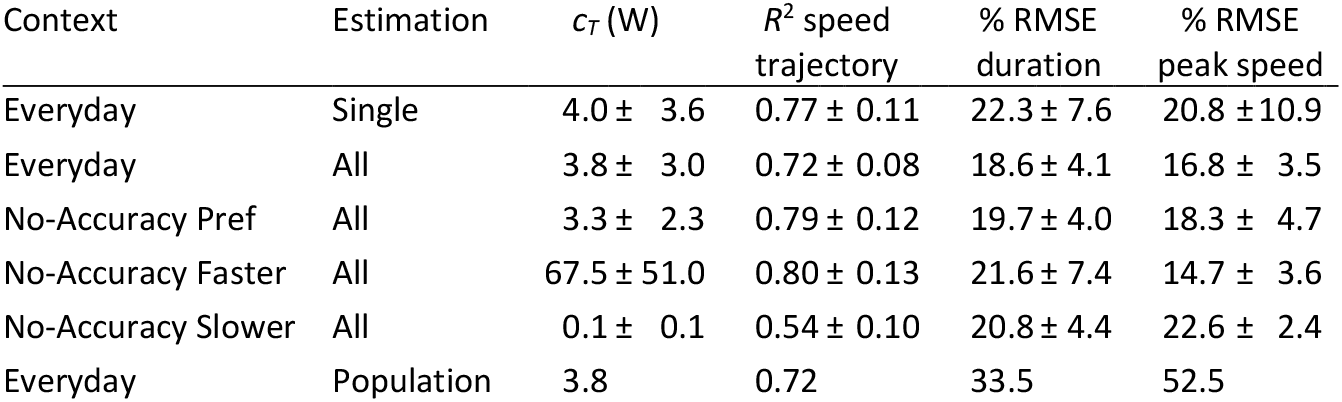
Experimental results for each reaching context, including Everyday reaches at self-selected speed and No-accuracy reaches under Preferred, Faster, and Slower contexts. Time valuation *c*_*T*_ and goodness-of-fit for optimal control speed trajectories (*R*^2^), power law durations *D* (% RMSE) and peak speeds *V* (% RMSE). Time valuation was estimated three ways: from Single observation per individual, All observations per individual, or Population-wide as a global parameter. Values are reported as mean ± SD across participants (*N* = 9).

| Context | Estimation | $c_T$ (W) | $R^2$ speed trajectory | % RMSE duration | % RMSE peak speed |
| --- | --- | --- | --- | --- | --- |
| Everyday | Single | $4.0 \pm 3.6$ | $0.77 \pm 0.11$ | $22.3 \pm 7.6$ | $20.8 \pm 10.9$ |
| Everyday | All | $3.8 \pm 3.0$ | $0.72 \pm 0.08$ | $18.6 \pm 4.1$ | $16.8 \pm 3.5$ |
| No-Accuracy Pref | All | $3.3 \pm 2.3$ | $0.79 \pm 0.12$ | $19.7 \pm 4.0$ | $18.3 \pm 4.7$ |
| No-Accuracy Faster | All | $67.5 \pm 51.0$ | $0.80 \pm 0.13$ | $21.6 \pm 7.4$ | $14.7 \pm 3.6$ |
| No-Accuracy Slower | All | $0.1 \pm 0.1$ | $0.54 \pm 0.10$ | $20.8 \pm 4.4$ | $22.6 \pm 2.4$ |
| Everyday | Population | 3.8 | 0.72 | 33.5 | 52.5 |

#### Everyday reaching predicted across distances, participants, and population

Further agreement was demonstrated across a larger range of distances. Here, each individual’s *c*_*T*_ was identified from all their data across up to fifteen distances. Experimental hand speed trajectories agreed well with bell-shaped predictions (Fig. 4, top row), and movement durations increased approximately with *D*^1/4^ and peak speeds with *D*^3/4^ (p = 3.3e-193), in agreement with the power law approximations and optimal control (see Table 1, Everyday All). A population-wide prediction was also nearly as accurate as individual-specific ones. The population-wide *c*_*T*_ was about the same, 3.8 W, and applied to all data yielded about 15% greater root-mean-square error (RMSE) for duration and 36% for peak speed (compare results for Population against All individual-specific, Table 1). Thus, most participants made Everyday movements at similar speeds.

**Fig. 4.**
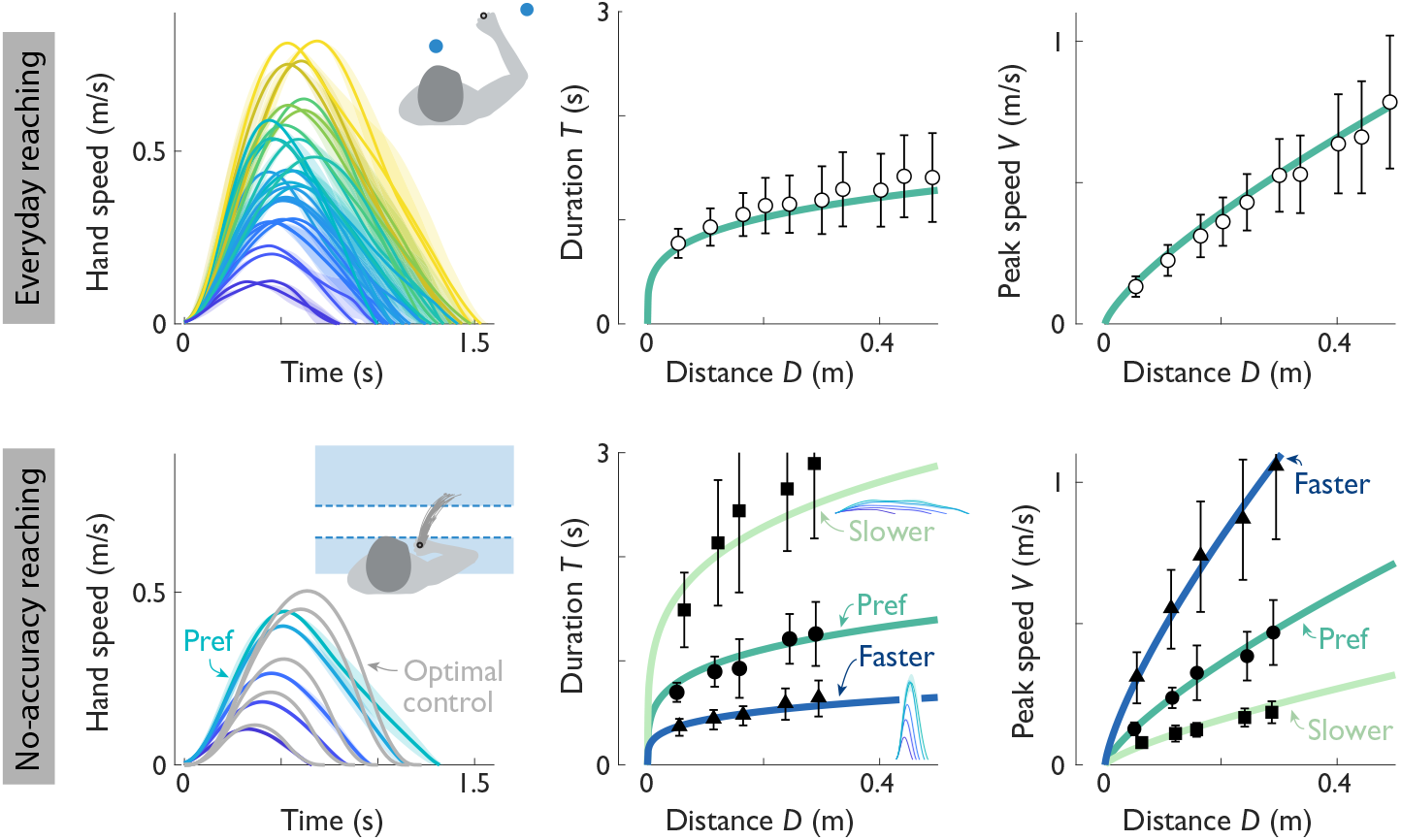
Experimental reaching data: (left to right:) speed trajectories, durations, and peak speeds across reaching distances *D*. (Top row:) Everyday reaches at preferred speed from all participants (*N* = 9, mean ± s.d. for distances 0.025 to 0.60 m), modeled using individualistic time valuation *c*_*T*_. (Bottom row:) No-accuracy reaches, where participants reached various distances between separated half-plane targets under three contexts: Preferred speed, Faster and Slower than preferred. Each context yielded a distinct *c*_*T*_. Insets show experimental hand speed trajectories for Faster and Slower contexts. Table 1 summarizes quantitative results across conditions.

#### No-Accuracy reaches predicted by Energy-Time

These reaches had little to do with maximizing accuracy (Fig. 4, bottom row). A separate set of “No Accuracy” reaches were conducted between large targets each about half the size of the workspace (two half-planes separated by varying distances). If the speed-accuracy trade-off were a determinant of Everyday reaches, then relaxing target size should result in faster preferred speeds. But this was not the case: No Accuracy reaches at preferred speed had similar bell-shaped trajectories, durations, and speeds as Everyday, and showed no significant differences in peak speeds and durations (p = 0.60 and 0.55, respectively for matched distances). When not under time or accuracy pressure, humans appear to prefer smooth, relaxed Everyday movements as predicted by Energy and Time.

#### Contextual modification of No Accuracy reaches

The No Accuracy reaches were also consistent within a subjective movement context. We tested this by changing the context via instructions for participants to reach “Faster” or “Slower” than preferred in No Accuracy conditions. The contexts caused experimental speeds to differ more than five-fold, but the movements were nevertheless consistent with the Energy-Time model (Fig. 3, bottom row). The Slower speed trajectories were lower in speed and longer in time, and the opposite for Faster reaches. The durations still increased approximately with *D*^1/4^ and peak speeds with *D*^3/4^ for both contexts, which differed only in time valuation.

If all movements are indeed optimal, they should agree with the same universal predictions. The predicted surfaces (Figs. 3B-C, Eqs. 2-3) should apply to any distance and, by selecting *c*_*T*_, any individual and context. We therefore compared data from all individuals and conditions (Everyday, plus Fast, Preferred, Slow for No Accuracy) with the surfaces (Fig. 5 left column shows same surfaces as predictions Figs. 3B & C), comprising average speeds differing two-fold across individuals (at preferred context) and five-fold across contexts. All reaches by an individual and context were represented well by the distance dependence predicted by a single *c*_*T*_ (solid lines in Fig. 5 left; middle column shows same data collapsed across *c*_*T*_; about 20% RMSE). Participants thus reached in highly consistent ways, regardless of distance, accuracy constraint, or subjective context.

**Fig. 5.**
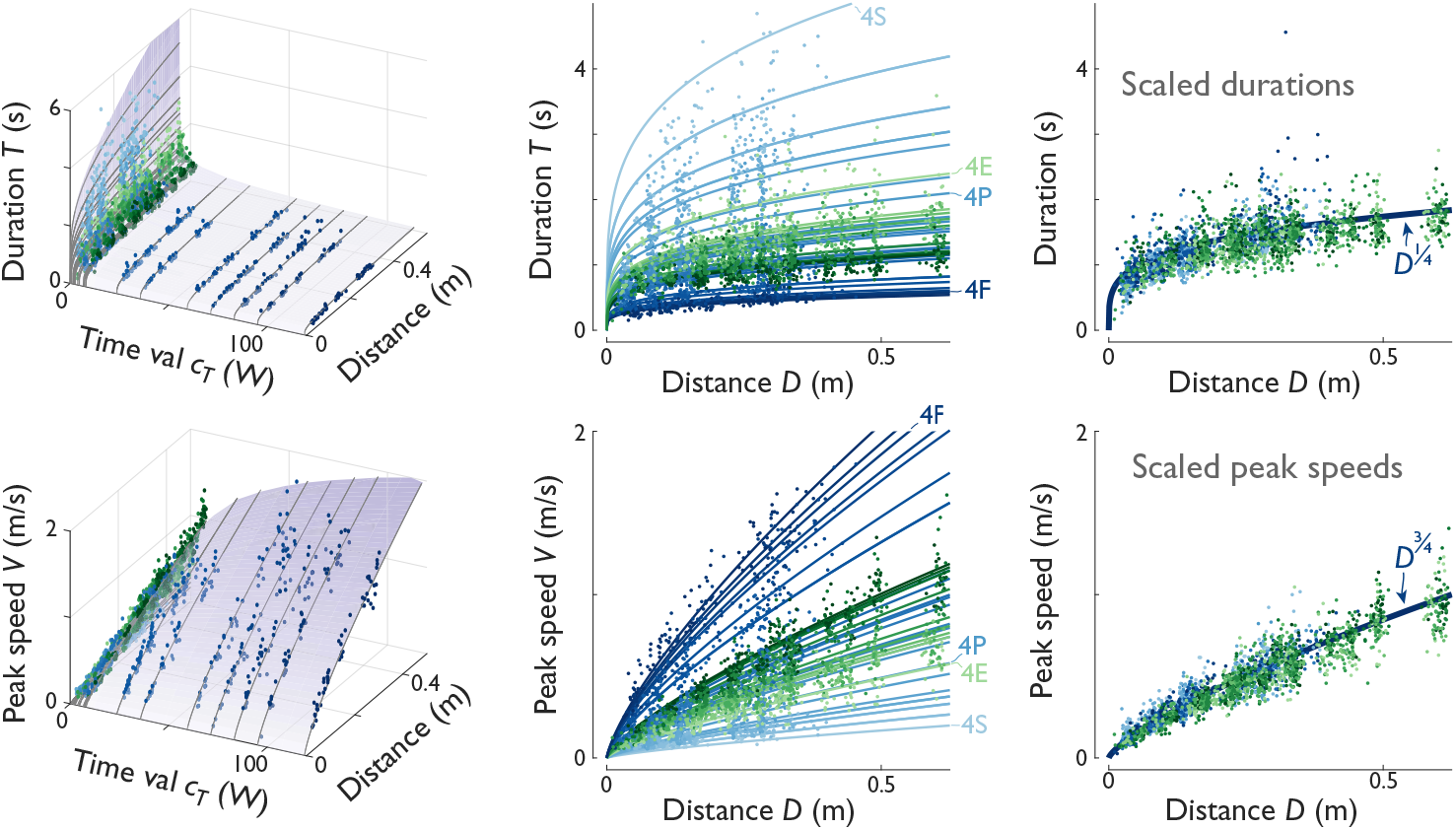
Human reaching durations (*T*, top row) and peak speeds (*V*, bottom row) across all distances, participants (*N* = 9), accuracies, and contexts. (Left column:) Duration and peak speed as functions of distance *D* and time valuation *c*_*T*_; (Middle:) Duration and peak speed versus distance, collapsed across all *c*_*T*_. (Right:) Normalized duration and peak speed versus distance, with generic time valuation (*c*_*T*_ = 1.6 W). Data show all conditions: Everyday and No-accuracy (Preferred, Faster, Slower; see Fig. 3). Individualized *c*_*T*_ values (solid lines, color-matched to data points) yield RMSEs of 20.2% for duration, 18.1% for peak speed. Normalized data (right column) show durations scaling with *D*^1/4^ and peak speeds with *D*^3/4^, reducing inter-subject and inter-context variability by 76% for duration (s.d. unnormalized 0.66 s, normalized 0.16 s) and 78% for peak speed (s.d. 0.27 m/s, 0.06 m/s). Representative reaches for participant 4 are labeled: Everyday (4E), Slow (4S), Preferred (4P), and Fast (4F).. Predictions from Eqs. 2 and 3 are shown as surfaces in left column, consistent with Figs. 3B & C.

#### Universality of Energy-Time predictions

The Energy-Time model also predicts that all reaches should scale similarly with distance (*D*^1/4^ for duration, *D*^3/4^ for peak speed), regardless of how fast. This was demonstrated by normalizing each individual’s time valuation to a generic one (Fig. 5 right). Specifically, the durations for each individual and context were multiplied by 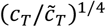, where *c*_*T*_ and 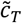 are valuations for an individual and for all averaged data, respectively; peak speeds were multiplied by the inverse. Normalization caused the variability across individuals and contexts to reduce by 76%, suggesting that most of the differences are due to the valuation of time. All reaches for all individuals and contexts therefore scaled with *D*^1/4^ for duration and *D*^3/4^ for peak speed. There was thus no qualitative difference in reaching regardless of distance, (modest to low) accuracy demand, and faster or slower context.

## Discussion

Understanding everyday human reaching movements requires a model that balances physiological constraints, subjective preferences, and task demands. We hypothesized that an Energy-Time objective could explain these movements by integrating metabolic energy expenditure with time valuation. Using a simple power law (Eqs. 2 and 3) with just one parameter (*c*_*T*_), the model successfully predicted reaching trajectories, durations, and peak speeds across different distances and contexts. This parameter was identifiable from a single experimental observation and generalized to other reaches by the same individual (Fig. 3). The model also accounted for movements with no accuracy demands, and with subjective faster and slower contexts, by adjusting time valuation alone. Despite its simplicity, the Energy-Time model outperformed prior models in its ability to explain a wide range of movements while remaining interpretable and physiologically grounded.

The proposed Energy-Time model addresses key limitations of prior models. Most previous approaches focus on either bell-shaped trajectories or movement durations but not both simultaneously. Accuracy has been the prevailing theory for fast, accurate movements but does not apply well to Everyday movements here, which were self-paced and undemanding of speed. After all, humans frequently reach for objects like cups or pens without fixating their gaze accurately or rushing to completion. Furthermore, the No-Accuracy reaches here were about the same speed as Everyday ones, whereas the speed-accuracy tradeoff would predict faster movements once relaxed. Moreover, most prior models fail to account for subjective context without introducing additional empirical parameters. In contrast, our model uses a single parameter *c*_*T*_ to predict movement trajectories, durations, and peak speeds after observing just one reach per individual and context.

The proposed model also predicts movements beyond the present experiment. Unsurprisingly, the model predicts that moving the arm with added mass should cost proportionally more energy (Eq. 1, treating *M* as total mass). Less obvious is its prediction of modest effects on preferred durations and slower peak speeds, changing only with *M*^1/4^ (Eqs. 2-3). These three predictions align with trends reported by Bruening et al. (20) for reaching with added arm mass (Fig. 6**A**-**C**). The metabolic power model (Eq. 1) has also been tested with separate cyclic reaching experiments (Fig. 1**C** (14)) and is consistent with independent data from Shadmehr et al. (12) on controlled reaching distances and durations (Fig. 6**D**). Although initially developed to explain force production costs during cyclic leg swinging (21) (Fig. 6**E**), the energy model applies broadly to other tasks such as ankle bouncing (22) and isometric quadriceps contractions (23) (Fig. 6**F**). These other data also test the energetic cost for performing mechanical work (see Methods), which is critical to movement in general (22, 24), and for predicting optimal control trajectories here (Fig. 3D). But the force-rate cost seems to dominate energetic cost (14) in everyday movements, as well as those with added arm mass.

**Fig. 6.**
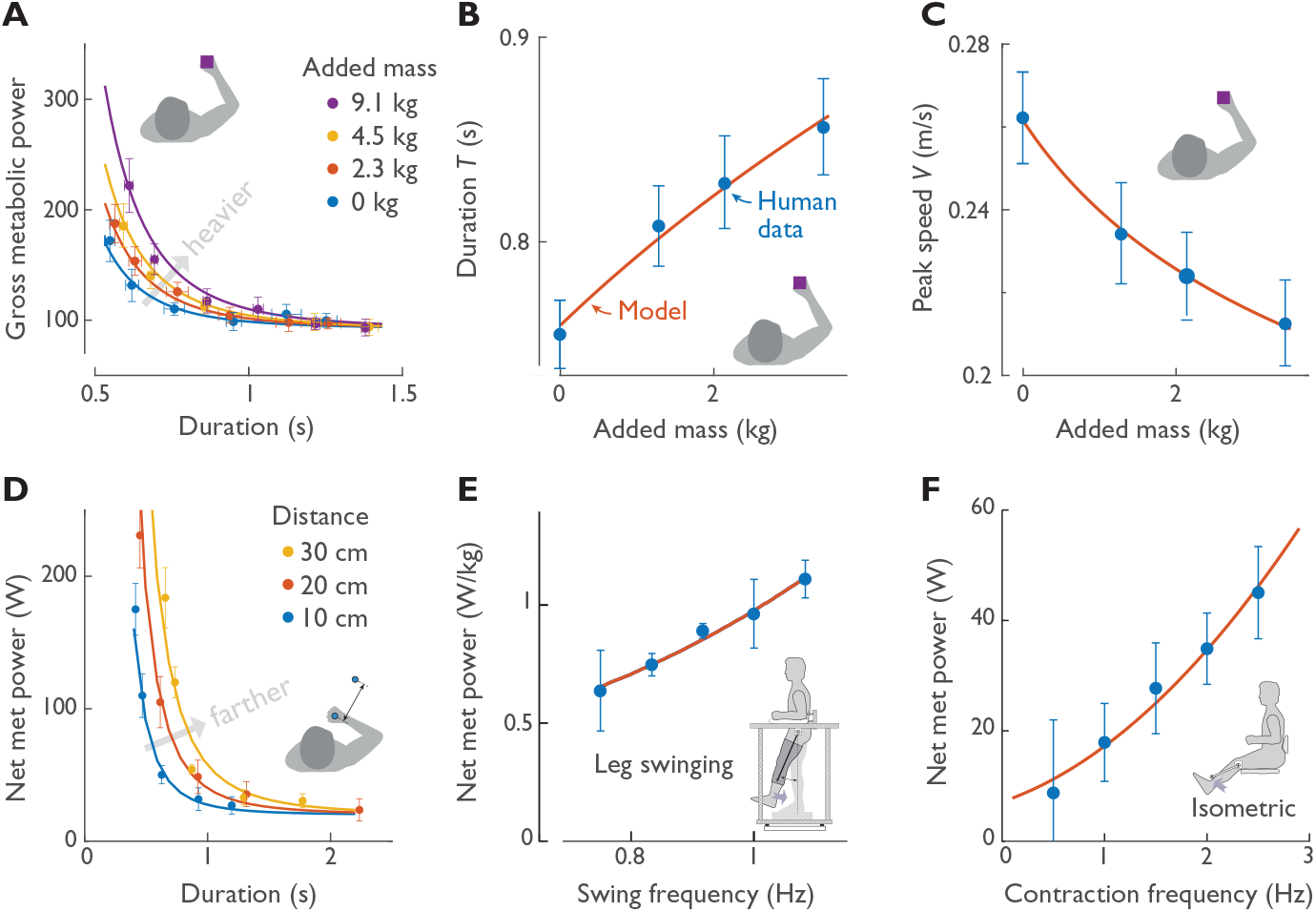
Energetic cost model compared with four independent experiments: (**A** - **C**) reaching with added arm mass, (**D**) reaching different distances, (**E**) cyclic leg swinging at different frequencies, and (**F**) cyclic isometric knee extension at different frequencies. (**A**) Metabolic power vs controlled duration with added arm mass, (**B**) preferred duration and (**C**) preferred peak speed versus added mass (all digitized data^30^), compared to model predictions (Eqs. 1-3). (**D**) Metabolic power vs. controlled duration for reaching of various distances (digitized data^5^), compared to model predictions. (**E**) Metabolic power versus controlled frequency of isolated leg swinging^31^ and (**F**) metabolic power vs. controlled frequency of cyclic isometric quadriceps contractions^32^. The present force-rate energetic cost model is shown for comparison (solid lines, **A**-**D**), with metabolic power prediction *MDT*^−4^ (Eq. 1, converting cost per contraction into cost per time *T*), *M*^1/4^*D*^1/4^ (Eq. 2) for preferred duration (**B**), and *M*^−1/4^*D*^3/4^ (Eq. 3) for peak speed (**C**). For lower extremity tasks, the same force-rate cost model predicts the metabolic costs of (**E**) leg swinging or (**F**) quadriceps contractions^32^ at different frequencies. For **A-D**, present model predictions are shown as solid lines, substituting for previously published empirical fits^5,30^.

The subjective valuation of time was remarkably stable for each individual and movement context. Individuals varied considerably in time valuation *c*_*T*_, as evidenced by standard deviations of movement speed comparable in magnitude to the means for all conditions (Table 1). But once identified, that quantity could nonetheless predict across all other reaching distances for that individual (Fig. 3). Moreover, reaches for all individuals and contexts were unified by a single relationship (Eq. 1 yields Eqs. 2 and 3, surfaces in Fig. 5 left column), despite elimination of accuracy demands and despite vague verbal instructions for Fast, Preferred, or Slow. This unification is further demonstrated by normalizing by *c*_*T*_, which reduced the inter-subject and inter-context variabilities in duration and peak speed by about three-quarters (Fig,. 5, right column). Although this was expected, we did not anticipate that all reaches for an individual would align closely along a single *c*_*T*_ value (solid lines in Fig. 5 left and middle). Thus, *c*_*T*_ appears sufficient to quantify highly individualistic interpretation of a movement context, and stable enough to predict all other movements within that context. We suspect that subjective contexts might well vary day-to-day due to factors like mood, food intake (25), or caffeine consumption (26), but the observed stability highlights the short-term robustness of *c*_*T*_ as a determinant of motion.

The Energy-Time model is highly testable because its parameters can be identified independently from present data. Most reaching models identify parameters through data fitting, but our optimal control parameters were independently derived from separate cyclic reaching experiments (14), making trajectory predictions (Fig. 3**D**) independent. (It would likely be more accurate to use data for discrete rather than cyclic reaching, but this highlights the model’s generalizability.) While duration and peak speed predictions (Eqs. 2–3) required fitting, they relied on as little as a single datum per individual (Fig. 3**E**). Previous models have included effort-like objectives such as jerk, torque change, or variance; metabolic energy extends this family but offers greater physiological relevance and independent falsifiability. Every major variable in the present model (metabolic energy, *M, D, T, V, c*_*T*_) has been experimentally tested here or elsewhere (Fig. 6).

Our model suggests a unified explanation for how bell-shaped speed profiles are produced across previous movement objectives. Four decades of hypotheses for minimizing jerk (9), torque change (3, 11), endpoint variance (4), and force rate share a common foundation in third-order dynamics, with second-order dynamics for arm motion states such as position and velocity, plus a third state for first-order activation dynamics (4) or equivalent. An instantaneous change in any third-order input produces a linear (ramp-like) initial increase in force or acceleration and a quadratic increase (bell-shaped leading edge) in velocity, and similarly for the trailing edge. Biological muscles have first-order dynamics at minimum (23), and can therefore impose at least equivalent constraints on speed changes. This illustrates a limitation of the inverse optimization approach of inferring neuromotor movement objectives from movement data, because many different models can produce similar bell-shaped speed profiles. The Energy objective is only one such model, but differs from other, seemingly-plausible objectives, by being physiologically measurable and testable.

There are several limitations to this model. First, the force-rate cost is supported better by data than theory. There is a long history of energy cost observations for fast bursts of muscle force, not explained by work (27, 28). Most musculoskeletal models include first-order dynamics for calcium activation (29, 30), which might seem like a suitable mechanism. But we believe a separate step prior to force production that we term “force facilitation” is slower and thus more rate-limiting, and better explains calcium pumping energy costs (23). But the mechanism is poorly understood, and so force-rate cost here is primarily a crude, first-order representation of energetics data.

Second, we presently lump subjective temporal effects into modulation of time valuation *c*_*T*_, which is not directly observable. Once determined, it is predictive of movements across distances and therefore testable, but how it is modulated by the central nervous system is unclear. Time has long been recognized as a factor in self-selected movements (31, 32), with individualistic valuation perhaps applying across tasks (6, 19). Most previous time models have been nonlinear, within for example a time-varying cost of time (7, 19) or temporal discounting of reward and energy (12). But we find a linear cost of time, in walking (33), reaching (14), and single joint movements (8) to be simpler yet still predictive. After all, basal or resting energy cost is linear in time. It seems reasonable to augment that literal cost with subjective modulation, interpretable as the additional energy one is willing to expend to save a unit of time. But we do not know how reward affects time valuation, because reward is often implicit and unobservable as well. We present the following challenge: Predict how quickly an individual will reach for a US$100 bill, given an observed duration for a reach to $1. We suspect the differences are small and difficult to predict with any present theory, including ours.

Another limitation is that the present model did not include accuracy. We envision the speed-accuracy tradeoff as an optimization constraint, which may be active during demanding tasks such as throwing a baseball fastball and thus dominate energy as a concern. But energy and accuracy are somewhat ambiguous to distinguish because both favor slower movements and predict similar bell-shaped trajectories. The ideal experiment would manipulate energy, accuracy, and time as simultaneous tradeoffs, for example in swatting a fly. A smaller and more nimble fly would presumably demand more accuracy or faster speed, and require more energy to swat. But we also believe the speed-accuracy constraint is inactive during many everyday movements, by strategy or by design. For example, one can thread a needle or target a computer mouse by first moving ballistically and inaccurately only at first, and then switching to non-ballistic visual feedback control for accuracy. In that case, the ballistic, minimum-variance hypothesis would not apply. Everyday tools such as cups, pens, and fly-swatters also seem designed to lessen the accuracy demand. Although accuracy can be an important movement constraint, we intentionally focused on Everyday, easy reaching because it is highly ecological.

The Energy-Time model provides a unified framework for understanding everyday human reaching movements by balancing energy expenditure, time valuation, and subjective factors. It grounds prior theories of minimizing jerk or variance in physiological principles such as metabolic cost and force-rate dynamics. Unlike previous models that rely on abstract or task-specific parameters, the Energy-Time model uses independent, physiologically measurable variables like energy expenditure and time, which are fundamentally important for all living organisms down to individual cells and neurons (17). It also compresses individual- and context-dependent factors into a single, testable trade-off parameter *c*_*T*_. Despite the limitations above, the model remains highly interpretable and testable across tasks and datasets alike. Future research could explore how reward modulates time valuation or how energy interacts with accuracy demands in more complex tasks. By integrating physiological constraints with subjective preferences into a cohesive framework, the Energy-Time model advances our understanding of human motor control while offering practical applications in behavior analysis, motor rehabilitation, and robotics.

## Methods

There were three main components to this study: a simple energy-time optimal control, a power law approximation to the model, and an experimental test. The optimal control model, detailed previously (14), is briefly summarized here. The task objective is to determine time-varying force or torque trajectories over duration *T* to

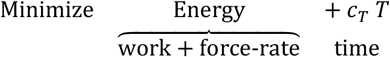

where the metabolic Energy term has hypothesized proportional components for work and force-rate. Muscles perform work on the arm with approximately proportional energetic cost (39) attributed to actin-myosin cross-bridge action (40). The force-rate cost is for rapidly producing force, because activation and deactivation of muscle is also costly. This term increases approximately with the force and inversely with its time duration, attributable to active transport of myoplasmic calcium (23, 41–43) and supported by metabolic cost data for a variety of movements (22–24). Energy cost per contraction is thus expected to increase approximately with the work performed and the amplitude of force-rate.

Optimal control is required to compute the full movement trajectories (14). The decision variables are joint torques τ_*i*_(*t*)(*i*=1 for shoulder and 2 for elbow) and final time *T*, for joint angle trajectories *θ*_*i*_. The energy terms are proportional to positive increments of the rates of joint mechanical work and force-rate,

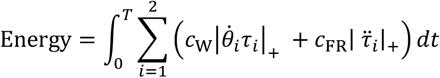

where |*x*|, = max(0, *x*)and *c*_+_ is the work coefficient, *c*_FR._ the force-rate coefficient.

These are subject to constraints for initial and final (target) positions, starting and ending at rest, with a two-segment model of arm dynamics including manipulandum inertia. These energy and time terms are needed for optimal control predictions (Fig. 2 and 3, optimal control), which included bell-shaped velocity profiles as well as movement durations *T* and peak speeds *V*.

Further analysis shows that the durations and peak speeds are also approximated by simple power laws. Proportionalities for the two energy components are

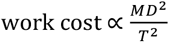

and

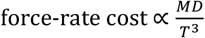

Although both contribute to overall energy cost, the force-rate cost should have greater influence on movement speed if reaching durations and distances are relatively small. We found that neglecting the work term and minimizing the remaining force-rate and time components yields far simpler power law predictions for *T* and *V* (Equations 2 and 3, respectively) that still agree well with optimal control (Figure 2D). Previous experiments also show that force-rate dominates the overall energy cost of cyclic reaching (14), especially at faster cycle frequencies. We therefore treated the power law approximations as a simplification of more complex optimal control predictions that facilitated comparison with experimental data. For example, the power laws suggest that data across individuals and contexts may be normalized by 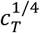 as another test of the model (Fig. 5).

The power law approximations were compared to optimal control as follows. Each approximation has a coefficient of proportionality, determined by least-squares fit to optimal control; the parameters *k*_*t*_ and *k*_ν_ that best fit simulated reaches with distance *D* spanning 0.01–0.5 m, and time valuations *c*_*T*_ spanning 0.02–120 W, were *k*_*t*_ = 2.26; *k*_ν_= 0.92. *R*^2^values for power-law approximations to optimal control durations *T* and peak speeds *V* were 0.980 and 0.983, respectively.

### Experiment

Healthy adult participants (N = 9) made reaching planar movements between visually-displayed targets with the right arm. The arm was supported by a two-joint robot manipulandum (KinArm), used passively to measure motions. Everyday targets were 2.5 cm diameter and separated by up to fifteen distances *D* of 2.5 – 60 cm. Participants were asked to start and move at self-selected time and speed and stop within the other target, with the target disappearing at that point. Movement duration *T* was defined to end when hand speed (defined along axis between targets) was reduced below 0.25 cm/s. Peak speed *V* was defined as the maximum axial hand speed in each trajectory. The relatively large target size and relaxed ending criterion (e.g., without a target hold period at zero speed) was intended to emulate everyday tasks such as grabbing a cup. There were five trials per distance (averaged together for one trajectory, duration, and speed per condition). Trials were initiated at fixed 5 s intervals indicated by an auditory cue, to avoid incentive to move faster to reduce overall experiment time. Prior to data collection, participants provided their written informed consent according to institutional review board procedures.

“No accuracy” reaches were defined as movements between two half-planes separated by distance *D*. The half-planes were oriented to approximately maximize their free workspaces, although participants were free to strike the manipulandum’s physical limits to arrest motion without affecting success (although nobody did), so that the targets were effectively of unlimited size. The No accuracy reaches were performed at self-selected (Preferred) speed, and Faster and Slower than Preferred. There were five distances ranging 2.5 – 30 cm, and five trials for each distance. The entire experiment entailed 150 reaches and took about 15 min.

## Data availability statement

The data and software for this study will be shared in a publicly accessible GitHub repository.

